# Viral delivery of compact CRISPR-Cas12f for *in vivo* gene editing applications

**DOI:** 10.1101/2024.02.06.578965

**Authors:** Allison Sharrar, Zuriah Meacham, Johanna Staples-Ager, Luisa Arake de Tacca, David Rabuka, Trevor Collingwood, Michael Schelle

## Abstract

Treating human genetic conditions *in vivo* requires efficient delivery of the CRISPR gene editing machinery to the affected cells and organs. The gene editing field has seen clinical advances with *ex vivo* therapies and with *in vivo* delivery to the liver using lipid nanoparticle technology. Adeno-associated virus (AAV) serotypes have been discovered and engineered to deliver genetic material to nearly every organ in the body. However, the large size of most CRISPR-Cas systems limits packaging into the viral genome and reduce drug development flexibility and manufacturing efficiency. Here, we demonstrate efficient CRISPR gene editing using a miniature CRISPR-Cas12f system with expanded genome targeting packaged into AAV particles. We identified efficient guides for four therapeutic gene targets and encoded the guides and the Cas12f nuclease into a single AAV. We then demonstrate editing in multiple cell lines, patient fibroblasts, and primary hepatocytes. We then screened the cells for off-target editing, demonstrating the safety of the therapeutics. These results represent an important step in applying *in vivo* CRISPR editing to diverse genetic sequences and organs in the body.

## Introduction

Clustered Regularly Interspaced Short Palindromic Repeats (CRISPR) and CRISPR-associated (Cas) systems are versatile tools for targeted gene editing.^1–6^ The clinical promise of CRISPR-Cas technology recently took a major step forward with the approval of Casgevy as an *ex vivo* gene editing treatment for sickle cell disease. Several clinical trials are underway using gene editing as a tool to create *ex vivo* therapies for hematologic diseases and cancers (CAR-T).^7^ Delivering the Cas editing machinery directly to patients through *in vivo* therapies is also advancing with clinical trials using lipid nanoparticles (LNP) to deliver editors to the liver.^8^ A major hurdle for LNP therapies is the lack of tissue distribution with most LNP formulations restricted to delivery to the liver.^9^

By contrast, natural and synthetic Adeno-associated virus (AAV) serotypes have been developed that target many tissues in the body, including the liver, lungs, eye, heart, muscles, kidneys, and the central nervous system (CNS).^10^ Several gene therapies using AAV have been approved by the FDA to treat diseases of the liver, eye, and CNS.^11^ Safe and effective AAV delivery of therapeutic gene editing cargo would therefore broaden the spectrum of accessible tissues.

The biggest limitation to delivering editing machinery by AAV is the relatively small genetic payload (4.5 kb).^12^ The most widely used CRISPR-Cas systems (Cas9 and Cas12a) are large proteins ranging from 120–160 kDa (3.2–4.3 kb) and require additional fusions like nuclear localization signals for effective editing in human cells.^5,13^ Since AAV deliver a genetic payload, the cargo must also include the requisite promoters and terminators for transcription and translation as well as separate promoters for the single guide RNA (sgRNA). Splitting the genetic cargo into multiple vectors or using minimal promoters with the smaller Cas nucleases are imperfect solutions to this problem and require significant tradeoffs in AAV packaging, genetic stability, payload expression, and drug development.^14,15^

The recent discovery of the highly compact CRISPR-Cas12f nuclease family (60 kDa) poses an elegant solution to delivery of editing machinery using AAV.^16–20^ The Cas12f family functions as a protein dimer around a single guide RNA, forming an editing complex that is equivalent to the larger Cas9 and Cas12a families but encoded in half the genetic space.^21^ We previously identified a novel set of Cas12f proteins, including AI1Cas12f1, and engineered them for editing in human cells.^20^ In this work, we developed AAV gene editing therapies encoding Al1Cas12f1 along with guides for several *in vivo* therapeutic targets and demonstrate editing and off-target analysis in cell lines and primary cells. This work is a critical step toward an effective and efficient AAV gene editing therapy.

## Materials and Methods

### Plasmid construction and AAV vector production

Nuclease and sgRNA plasmids used in transfection experiments were constructed as described previously.^20^ For AAV production, a codon-optimized Al1Cas12f1 gene was synthesized with an N-terminal SV40 nuclear localization sequence (NLS) and C-terminal nucleoplasmin NLS followed by a 3x human influenza hemagglutinin (HA) tag (Twist Biosciences). The Al1Cas12f1 sequence was inserted downstream of a CMV promotor in sgRNA expression plasmids containing inverted terminal repeat (ITR) sequences. AAV vectors were produced by BPS Bioscience, yielding 1.1-3.6×10^12^ viral genomes (vg) per mL.

### Cell culture and plasmid transfection

HEK293T and NIH3T3 cells were maintained in DMEM supplemented with 10% fetal bovine serum (FBS) and 1% penicillin-streptomycin. SMA patient fibroblasts were maintained in EMEM supplemented with 10% FBS, 1% penicillin-streptomycin, and 1X GlutaMAX (ThermoFisher). Mouse primary hepatocytes (Cell Biologics C57-6224F) were thawed in rodent hepatocyte thawing medium (Lonza MCRT50), plated with hepatocyte plating medium (Lonza MP100), and maintained in hepatocyte maintenance medium (Lonza MM250) according to manufacturer’s recommendations. All cells were kept at 37°C in an incubator with 5% CO2. Nuclease and sgRNA plasmids were co-transfected into HEK293T cells using Mirus Transit X2 reagent. Experiments were conducted in 96-well plates seeded with 5×10^4^ cells/well and transfected with 100 ng nuclease expression vector and 100 ng sgRNA expression vector according to manufacturer’s recommendations. Samples were incubated for 72 h.

### Al1Cas12f1-AAV-DJ transduction

Cells were seeded at 2-3×10^4^ cells/well into 96-well plates 24 h prior to AAV transduction at 100k MOI. HEK293T, mouse primary hepatocyte, and NIH3T3 experiments were incubated for 4-7 days with media changes every 3 days. Mouse primary hepatocyte media was exchanged every day according to Lonza recommendations. SMA patient fibroblasts were treated with 60 uM etoposide 5 h after seeding. Etoposide-containing media was removed at 24 h, immediately prior to AAV transduction. The SMA patient fibroblast experiment was incubated for 13 days and media was changed every 3 days.

### Genomic DNA extraction and sequencing

Genomic DNA was harvested with QuickExtract (LGC Biosearch Technologies) and amplified using KAPA HiFi polymerase with genomic site-specific primers. Amplicons from plasmid transfected samples were submitted for Sanger sequencing and editing efficiency was determined with TIDE analysis.^22^ Amplicons from AAV transduced samples were sequenced via Illumina iSeq with 150 bp paired-end reads and indel frequency was quantified with CRISPResso2.^23^ All editing data was plotted using Prism software.

### Off-target prediction

A custom version of Cas-OFFinder (ref) was used to search for off-targets related to PRSS1-g11 and SMN2-g1 guides in the human genome and PCSK9-g1 in the mouse genome. A DTTR (D = A, G, or T; R = A or G) PAM was used for Al1Cas12f1.^20^ For PRSS1-g11 and PCSK9-g1, all predicted off-targets where the combined DNA/RNA bulge size and number of mismatches totaled 4 or less were selected. For SMN2-g1, all predicted off-targets without bulges and <4 mismatches or <3 bulge size with one mismatch with were selected. There were many predicted off-targets with 2 mismatches and a bulge size of 1, so two were selected that had preferred PAMs. Off-target sites were amplified and sequenced via Illumina iSeq as described above.

## Results

### Al1Cas12f1 edits therapeutic targets via plasmid transfection

The *PRSS1* gene encodes cationic trypsinogen and is dysregulated in some patients suffering from hereditary pancreatitis.^24^ We designed 12 guides for Al1Cas12f1 targeting the exons of *PRSS1* to disrupt trypsinogen production (**Fig. 1A**). Single guide RNA (sgRNA) expression vectors were co-transfected with an Al1Cas12f1 expression vector and assessed for *PRSS1* editing in HEK293T cells. All guides exhibited *PRSS1* editing, with *PRSS1*-g11 having the highest efficiency (**Fig. 1B**). *PRSS1*-g11 targets exon 5 and has an ATTA PAM which is preferred by Al1Cas12f1.^20^

**Fig. 1.**
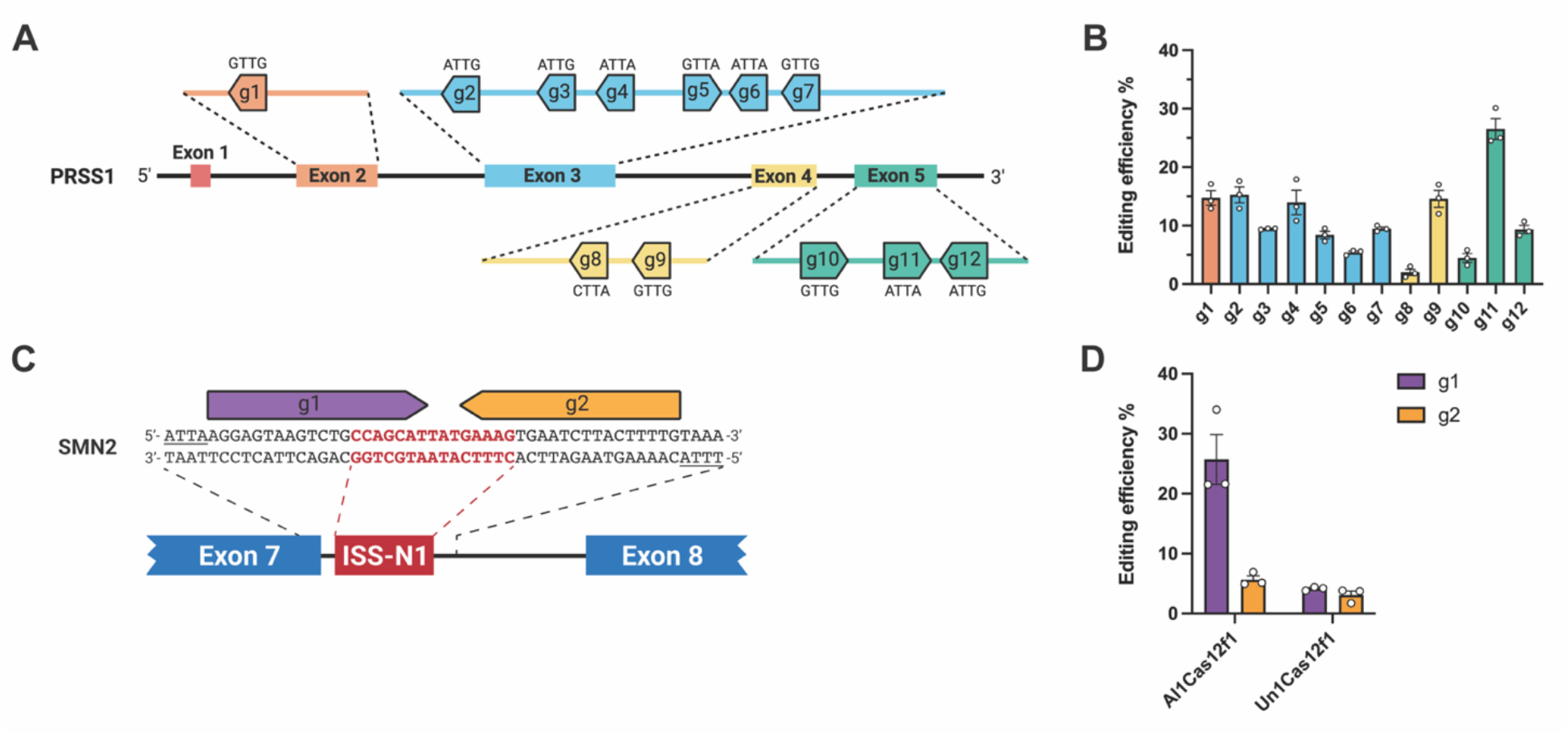
Guide selection for *PRSS1* and *SMN2*. **(A)** Locations and 5’ PAM sequences of *PRSS1* exonic guides (g1-12). **(B)** Editing efficiency of *PRSS1* g1-12 with Al1Cas12f1. **(C)** Locations of *SMN2* guides (g1-2) with cut sites in the ISS-N1 region. 5’ PAMs underlined. **(D)** Editing efficiency of *SMN2* g1-2 with Al1Cas12f1 or Un1Cas12f1. Editing performed in HEK293T cells using transient plasmid transfection and determined through TIDE analysis of Sanger sequencing reads. Bars represent mean ± SEM, n = 3 biological replicates.

Disruption of the ISS-N1 region downstream of *SMN2* exon 7 is a proven therapeutic technique for treatment of spinal muscular atrophy (SMA).^25^ We designed two guides with cut sites in this 15 nt long ISS-N1 region (**Fig. 1C**). Guides were tested for editing with Al1Cas12f1 and Un1Cas12f1^18^ (**Fig. 1D**). Un1Cas12f1 was unable to efficiently edit *SMN2* with either guide. By contrast, Al1Cas12f1 was capable of effective editing with *SMN2*-g1. The higher editing with Al1Cas12f1 on *SMN2*-g1 is likely due to the preference for ATTA PAM sequences whereas Un1Cas12f1 is unable to edit targets with ATTA PAM sequences.^20^

Al1Cas12f1 efficiently edits therapeutic targets in human and mouse cells via AAV delivery Al1Cas12f1 was packaged into AAV-DJ vectors^26^ with the top sgRNAs targeting human genes *PRSS1* and *SMN2*, as well as mouse genes *PCSK9* and *TTR* (**Fig. 2A**). *PCSK9* and *TTR* are therapeutic liver targets for treating familial hypercholesterolemia and ATTR amyloidosis, respectively.^8,27^ *PRSS1* and *SMN2* targeting vectors were transduced into HEK293T cells, yielding ∼40% editing efficiency (**Fig. 2B**). Additionally, the *SMN2* targeting construct was transduced into fibroblast cells from patients with SMA, yielding ∼10% editing efficiency. For the mouse targets, efficient editing could not be observed through plasmid transfection (data not shown), so multiple guides for *PCSK9* and one for

**Fig. 2.**
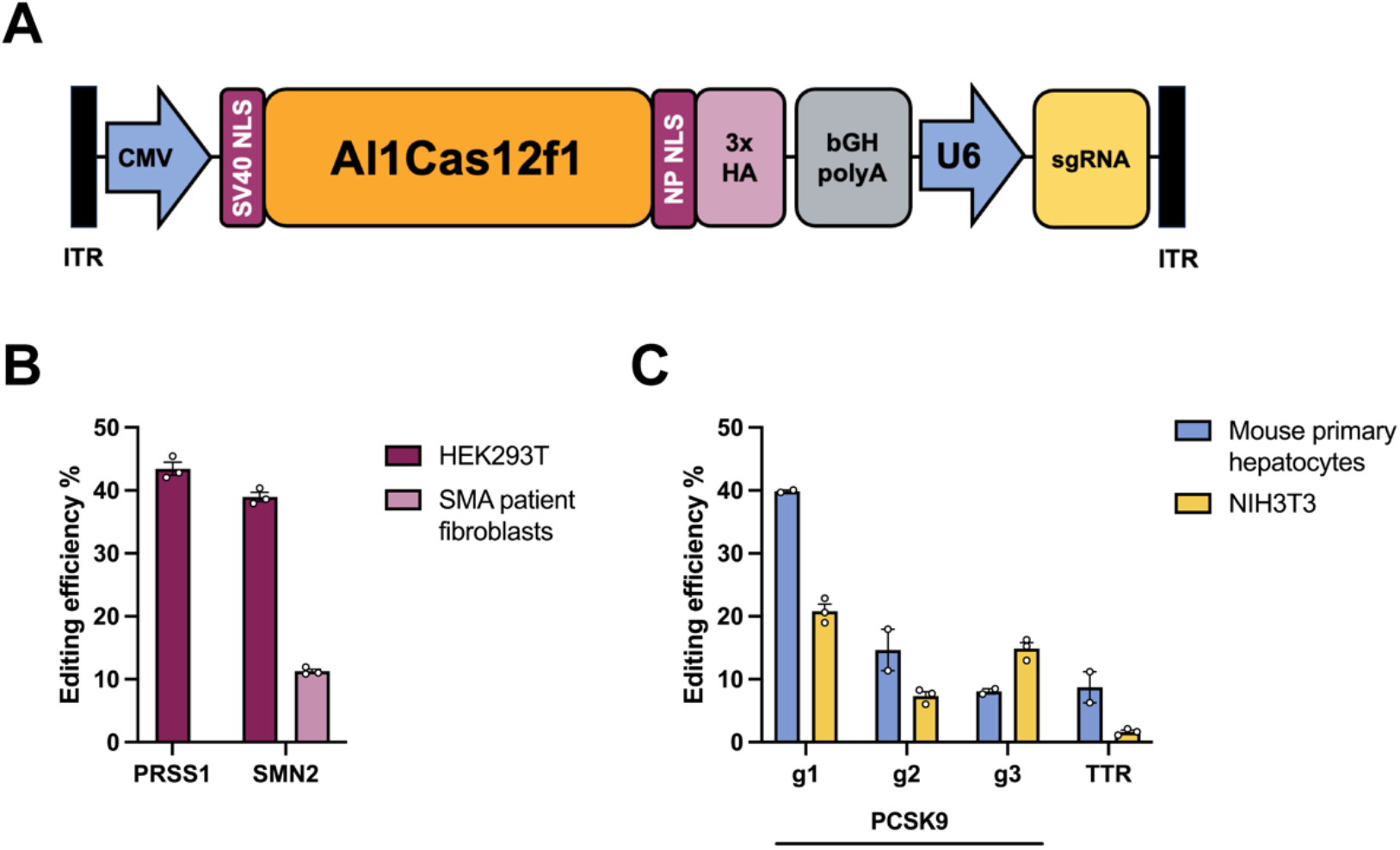
Al1Cas12f1 editing via AAV transduction. **(A)** Schematic diagram of Al1Cas12f1-AAV-DJ vector. **(B)** Editing efficiency at targeted sites in human cells with guides *PRSS1*-g11 or *SMN2*-g1 following AAV transduction. **(C)** Editing efficiency at targeted sites in mouse cells with guides targeting *PCSK9* or *TTR* following AAV transduction. Editing determined by analysis of NGS reads. Bars represent mean ± SEM, n = 2 (hepatocytes) or 3 (NIH3T3) biological replicates.

*TTR* were packaged directly into AAV for testing. *PCSK9* and *TTR* targeting vectors were transduced into NIH3T3 mouse cells and mouse primary hepatocytes. A maximum of ∼40% editing efficiency was reached for *PCSK9* in mouse primary hepatocytes, while *TTR* editing reached ∼10% (**Fig. 2C**).

### Low off-target editing with Al1Cas12f1-AAV-DJ

To assess the specificity of our novel Cas12f AAV construct, predicted off-targets were tested for top *PRSS1, SMN2*, and *PCSK9* guides. Off-target sites with the fewest mismatches and DNA/RNA bulges were amplified and sequenced from samples of the highest edited cell type. For *PRSS1*-targeting AAV, the highest off-target editing was below 2% for a two mismatch off-target site (**Fig. 3A**). For *SMN2*-targeting AAV, the highest off-target editing was below 1% for an off-target with one mismatch and a 2bp RNA bulge (**Fig. 3B**). For *PCSK9* in mouse primary hepatocytes, the highest off-target was under 0.1% editing efficiency, with two mismatches and a 1bp RNA bulge (**Fig. 3C**).

**Fig. 3.**
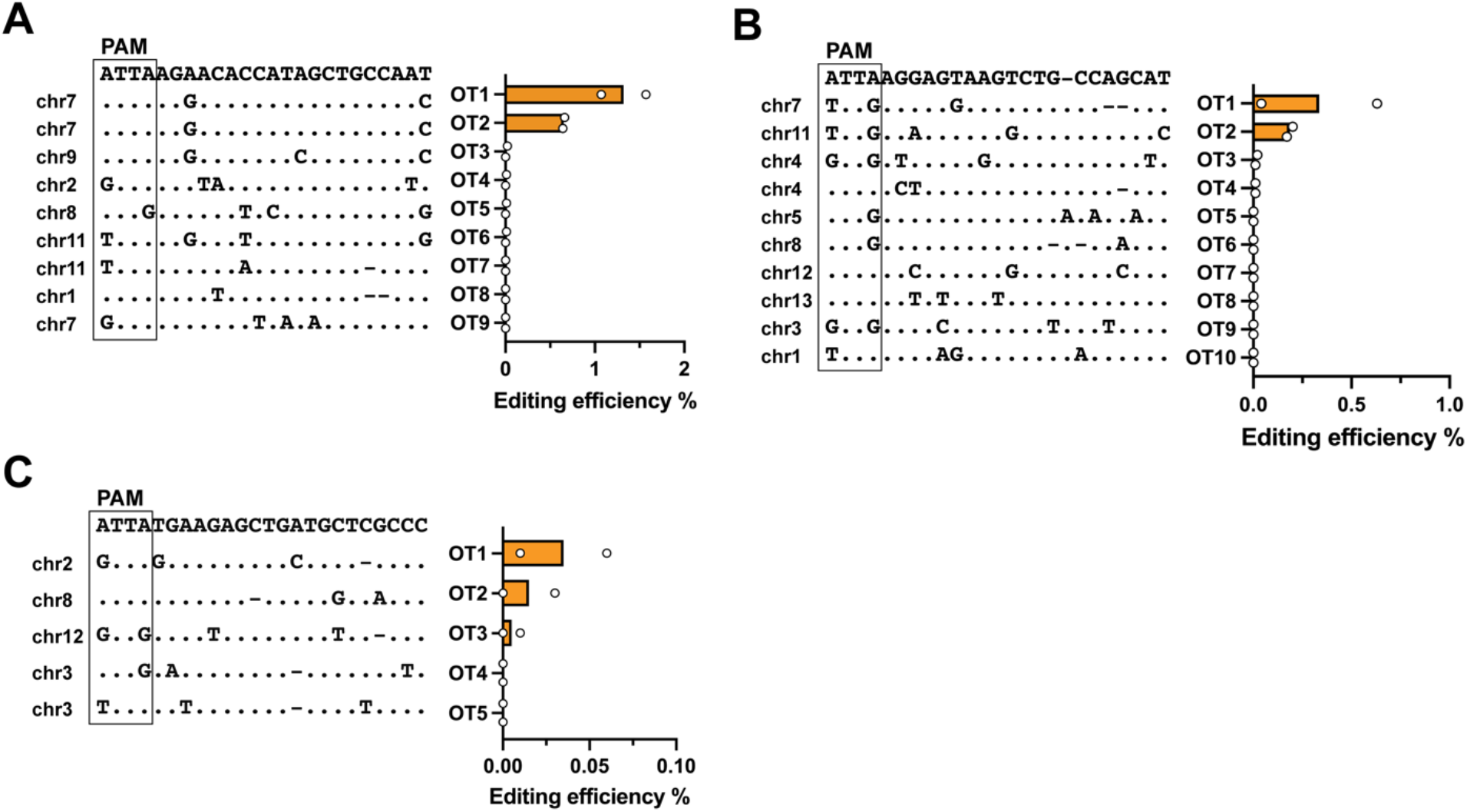
Editing of predicted off-target sites. Off-target sites were predicted *in silico* for **(A)** *PRSS1*-g11 in HEK293T cells, **(B)** *SMN2*-g1 in HEK293T cells, and **(C)** *PCSK9*-g1 in mouse primary hepatocytes. Editing efficiency was evaluated after AAV transduction and analyzed by NGS. Bars represent mean of two biological replicates.

## Discussion

The approval of the first CRISPR gene editing therapy highlights the tremendous potential for the technology to benefit human health. Advancing CRISPR technology to treat patients *in vivo* requires optimizing the delivery of the CRISPR-Cas editing machinery to patient tissues. AAV particles are a proven *in vivo* delivery technology used in multiple FDA-approved gene therapies.^10^ A major hurdle for AAV therapies is the physical limit for viral genome size. DNA payloads that are near this limit have decreased packaging efficiency and result in empty AAV particles.^28^ This precludes efficient packaging of many CRISPR-Cas systems like Cas9 or Cas12a. The CRISPR-Cas12f family is unique due to their very small size, roughly half of most CRISPR-Cas nucleases. Despite their small size, the enzyme are fully active editors which function using a dimeric structure that is based around a single guide RNA. The compact size of Cas12f nucleases allows for the inclusion of full-length promoters, guides, and other genetic elements that increase the efficiency and efficacy of AAV therapies.

In this work, we take advantage of the small size and broad PAM recognition of the recently discovered Al1Cas12f1 enzyme^20^ to package complete editing vectors in AAV particles. We first selected therapeutic gene targets (*PCSK9, TTR, SMN2*, and *PRSS1*) which are addressable *in vivo* using AAV delivery. We then screened multiple guides for each gene using plasmid transfection of HEK293T or NIH3T3 cells to identify the guides with the highest editing potential. The best performing guides for each of the gene targets contained an ATTA PAM, the preferred PAM for Al1Cas12f1^20^ and largely uneditable by other Cas12f systems.

We then designed AAV payloads encoding the complete Al1Cas12f1 sequence driven by the full-length CMV promoter and sgRNA driven by the U6 promoter. The payloads were packaged into the AAV-DJ serotype and evaluated for therapeutic gene editing in human or mouse cells. The *SMN2* and *PRSS1*-targeting AAV constructs efficiently edited human HEK293T cells and primary SMA patient fibroblasts (*SNM2* only). AAV targeting *PCSK9* and *TTR* effectively edited mouse primary hepatocytes as well as the mouse NIH3T3 cell line. These results demonstrate both the editing efficiency of Al1Cas12f1 and the ability of the nuclease and guide to be efficiently packaged in AAV particles. Off-target analysis of edited cells demonstrates the overall safety of the Al1Cas12f1 enzyme with most off-target sites showing very low levels of editing. Together, these data demonstrate the efficiency and utility of Al1Cas12f1 for *in vivo* gene editing applications.

## Acknowledgments

We would like to thank Prof. Joseph Bondy-Denomy for critical reading of the article.

## Authors’ Contributions

A.S and L.A.T. contributed to the study design, experiments, data analysis, and article writing. Z.M and J.S-A. contributed to experiments and data analysis, T.C. and D.R. contributed to study design. M.S. contributed to study design, data analysis, and article writing.

